# A Descriptive Analysis of *Streptococcus suis*-Associated Disease in Irish Pigs from 2010 to 2024: Serotypes, Pathology, and Antimicrobial Resistance

**DOI:** 10.1101/2025.08.17.670728

**Authors:** John Moriarty, Emmanuel Kuffour Osei, Colm Brady, Sara Salgado, Sebastian Alessandro Mignacca, Fiona Kane, Dayle Johnston, Laura Scanlon, Hannah Flynn, Marius Beumer, Rachel Reid, Philip Curran, Jennifer Mahony, Aine O’Doherty, Shane McGettrick, Cosme Sánchez-Miguel, Margaret Wilson, John G. Kenny

## Abstract

*Streptococcus suis* is a major cause of respiratory and systemic diseases in post-weaned pigs, leading to significant production losses and animal welfare concerns. This study provides the first long-term national level analysis of *Streptococcus suis*-associated disease (SSAD) in the Republic of Ireland. We examined the pig diagnostic submissions, characterised serotype distribution, antimicrobial susceptibility, and co-infection patterns from 2010 to 2024. The findings confirm that serotypes 9 and 2 or 1/2 were most frequently associated with disease. We observed a significant shift in recent years where serotype 9 has surpassed serotype 2 or 1/2 in number of occurrences. *S. suis* was frequently co-detected with viral pathogens including porcine reproductive and respiratory syndrome virus (PRRSV), porcine circovirus type 2, and swine influenza virus (SIV), as well as bacterial pathogens such as *Actinobacillus pleuropneumonia* and *Pasteurella multocida*, typically from pneumonic lungs. While resistance to tetracycline and erythromycin was high (44.4% to 65.8%), isolates remained susceptible to first-line beta-lactam antibiotics such as penicillin (7.9% resistance), ampicillin (5.5% resistance) and amoxycillin/clavulanate (0% resistance). The observed heterogeneity between and within herds challenges successful implementation of vaccination and highlights the need for ongoing disease monitoring. These findings provide the first in-depth assessment of SSAD in Ireland’s pig population which will offer valuable insights for future surveillance efforts, including genomic studies and supporting evidence-based strategies and vaccine selection for controlling *S. suis* in Irish pig sector.

## Introduction

*Streptococcus suis* is an important pathogen of intensive pig production industries globally, including Ireland. It represents a major cause of systemic disease and mortality in post-weaned pigs aged 5 to 10 weeks, imposing significant economic losses and animal welfare concerns [1]. In Ireland, the commercial pig sector is the third largest livestock industry, which mainly consists of integrated farrow-to-finish production systems, comprising approximately 300 commercial herds with approximately 140,000 sows and an annual output of 4 million pigs [2,3]. The pathogenic spectrum of *S. suis* ranges from subclinical carrier states to acute clinical infections and sudden death [4]. Early colonisation of the palatine tonsils and nasopharynx by *S. suis* results in an asymptomatic carrier state that serves as a reservoir for infection and facilitates the emergence of dominant pathogenic serotypes on farms. This is particularly evident during periods of stress including weaning, coinfections with respiratory pathogens, or exposure to adverse environmental factors such as high humidity and elevated CO_2_ levels in nursery units [5,6]. The tonsils are considered the primary route of entry for the pathogen, and breaching the mucosal barrier triggers a pronounced pro-inflammatory response leading to septicaemia, which can manifest as meningitis, polyserositis, pneumonia, arthritis, and acute deaths in weaned and less commonly, suckler pigs [7,8].

*Streptococcus suis* is an early coloniser of the upper respiratory tract and tonsils in new born pigs [9] and transmission occurs via aerosol and vertically from sow to piglet during farrowing [10,11]. Up to 100% of pigs in a herd may carry one or more serotypes, although in closed populations, there is typically a tendency for one strain to establish dominance and cause disease [12]. The virulence of clinical and subclinical isolates can differ substantially even within the same serotype, and emerging data suggest serotype switching is not uncommon [13,14]. Piglets from sows previously exposed to *S. suis* or of high parity tend to exhibit lower disease rates, likely due to transfer of maternal immunity [6]. A presumptive diagnosis based on clinical signs is confirmed by pathological examination and pathogen isolation, which establishes the aetiology and pathogenicity of *S. suis* isolates, thus enabling serotype characterisation from affected pigs [15]. At least 29 *S. suis* serotypes are confirmed to be true *S. suis* species based PCR typing and whole genome analysis [16]. Determining serotype of isolates from *S. suis*-associated disease (SSAD) affected pigs enables targeted vaccination using commercial or autogenous vaccines specific to disease-causing serotypes. However, the emergence of new pathogenic serotypes on farms presents an ongoing disease management challenge on the farm. There are marked geographical differences regarding serotypes and their disease manifestation. Although serotype 2 is reported as the predominant serotype in many countries globally, serotype 9 is emerging as the principal serotype in some European countries such as, Spain, Germany and The Netherlands [17,18]. In North America, serotypes 1/2, 1 to 9 and 14 are commonly associated with diseases, while serotypes 2, 1/2, and 3 predominates Australia; 2, 1/2, 3, and 4 in Asia, and 2, 1/2, 3, and 6 in South America [17,19,20].

Concurrent infection with other pathogens is common and significantly influence *S. suis* pathogenesis and disease severity. Synergistic interactions with bacterial agents such as *Mycoplasma hypopneumoniae* and *Pasteurella multocida*, as well as respiratory viruses including porcine reproductive and respiratory syndrome virus (PRRSV), porcine circovirus type 2 (PCV 2), and swine influenza virus (SIV), often exacerbate the risk and severity of pneumonia [21,22].

The lack of a universal vaccine that offers cross-protection against prevalent serotypes has resulted in widespread antibiotic usage [23]. Consequently, the emergence and dissemination of antimicrobial resistance (AMR) due to the selective pressure from antibiotic usage is well established and varies globally depending on usage patterns and implementation of national regulatory policies. For instance, tetracycline resistance in *S. suis* is considerably lower in Sweden than in other countries such as Denmark (52%) and Spain (95%), which is likely because usage of antibiotic growth promoters in animal feed has been banned in Sweden since 1986 [24,25].

This study is the first comprehensive report of SSAD in pigs in Ireland from 2010 to 2024. We document the occurrences of SSAD cases diagnosed through post-mortem examinations performed at the Veterinary Laboratory Service (VLS) of the Department of Agriculture, Food and the Marine (DAFM). We analyse *S. suis* serotype distribution, pathological lesions observed, and antimicrobial resistance profiles of the recovered isolates. Understanding the pathogenic serotypes circulating in the Irish pig population is vital for enhancing existing diagnostic protocols and implementing targeted-farm level disease prevention strategies. This baseline data will inform future surveillance programmes including genomic studies and guide evidence-based approaches to *S. suis* control in Irish pig production industries.

## Materials and Methods

### Case selection and data collection

We reviewed all pig carcass submissions to the VLS of DAFM from 1st January 2010 to December 2024 that received a diagnosis of SSAD. The submissions were made by specialist pig veterinary practitioners on behalf of herdowners and were typically ad hoc, with each submission usually including one or more pig carcasses for post-mortem examination. Non-carcass diagnostic submissions, such as swabs and organs alone, were excluded from this analysis unless otherwise stated. DAFM has recorded data of laboratory submissions on the laboratory information management system (LIMS) since 2003. We retrieved metadata for each case including herd of origin, pig age, clinical history, necropsy findings, ancillary laboratory test results, and final diagnosis with cause of death recorded by the pathologist. The full narrative post-mortem reports were reviewed to confirm lesions and diagnoses.

### Post-mortem examination and bacterial isolation

Standard post-mortem examinations were conducted by veterinary pathologists for diagnostic purposes. Tissues were selectively collected by the pathologist for further investigation based on clinical history and gross pathological observations at necropsy. For histopathological analysis, tissues were fixed with 10% neutral buffered formalin, embedded in paraffin, sectioned, and stained with haematoxylin and eosin (H&E) according to standard protocols. Furthermore, selected tissues were processed according to standard laboratory protocols described below.

Primary cultures were performed on sheep blood agar and MacConkey agar as part of routine bacteriological testing. Additionally, culture was performed on sheep blood agar and chocolate agar incubated in 7.5% CO_2_ at 37 ^°^C for 48 hours as this is often necessary for optimal colony growth. Following aerobic incubation at 37 °C, *S. suis* colonies typically exhibit alpha-haemolysis after 24 hours and beta-haemolysis after 48 hours on sheep blood agar. A single alpha-haemolytic colony was picked from sheep blood agar after 24 hours of incubation and subcultured onto fresh sheep blood agar to obtain pure isolate. Presumptive identification was carried out using standard microbiological methods including Gram staining (gram-positive cocci), catalase testing (negative), and oxidase testing (negative). Isolates were further characterised biochemically using the API identification system (bioMérieux, Marcy-l’Étoile, France) and confirmed by matrix-assisted laser desorption/ionisation time-of-flight mass spectrometry (MALDI-TOF MS) (Bruker Daltonik GmBH, Bremen, Germany), according to manufacturer’s instructions, prior to serotyping. Isolates were stored at –80 °C on cryo-protective beads. Each submission was associated with a unique submission diagnostic group (SDG) number in LIMS, which was used to trace archived isolates and link them to herd and case details.

During necropsy, tissues were collected at the pathologists’ discretion for virological testing. Viral pathogens detection was carried out by the Virology Division of the DAFM laboratories using PCR-based assays. For PRRSV, an in-house RT-PCR method based on Kleiboeker *et al.* [26] was employed. PCV2 detection was based on published protocols described by Brunborg *et al.* [27] and Zhao *et al.* [28]. Lastly, SIV detection was based in European Union Reference Laboratory (EURL) standard methods and Spackman *et al.* [29] for influenza A virus. In addition to virological testing, at the pathologists’ discretion, lung samples with pneumonic lesions were submitted to the DAFM testing laboratory in Cork for the detection of specific bacterial pathogens. PCR assays were performed for *Glaesserella parasuis* (formerly known as *Haemophilus parasuis*) based on McDowall *et al.* [30]; a duplex PCR for *Actinobacillus pleuropneumoniae* (APP) following Tobias *et al.* [31]; and *Mycoplasma hyopneumoniae* based on Marois *et al.* [32].

### Molecular serotyping

*S. suis* isolates were serotyped using a two-step multiplex PCR as described by Okura *et al*. [33]. This method is based on the capsular polysaccharide synthesis (*cps*) genes which are organised within a single locus on the *S. suis* chromosome. Isolates were cultivated overnight in Todd Hewitt broth at 37 °C for 18 hours under microaerophilic conditions using sealed jars and gas sachet (Anaerocult A, Merck, Germany). Genomic DNA was extracted from the overnight cultures using Monarch Genomic DNA Purification Kit (New England Biolabs, #T3010L) according to the manufacturer’s instructions. The PCR protocol consisted of two sequential steps. The first PCR (grouping) categorises isolates into *cps* groups by targeting genes conserved across multiple serotypes. Based on the grouping, the second PCR (typing) assigns an isolate to a specific serotype by detecting genes unique to that serotype within the assigned group. Primer sequences used in this study (Table S1) were reconstituted to 100 pmol/μl and then diluted to obtain 10 μM working stock. All PCR assays were performed using PCR Master Mix (2X) (Thermo Scientific, #K0171). The cycling conditions were as follows: an initial denaturation at 95 °C for 3 minutes, followed by 30 cycles of denaturation at 95 °C for 30 seconds, annealing at either 60 °C (grouping) or 58 °C (typing) for 90 seconds, and extension at 72 °C for 90 seconds (1 minute per kb); followed by final extension at 72 °C for 10 minutes. PCR products were subjected to agarose gel (1.5 %) electrophoresis at 95 volts for 45 minutes. Bands were visualised and photographed using GelDoc Go System (Bio-Rad Laboratories, USA). A subset of isolates was serotyped by SAN Group Biotech (Germany) using their proprietary PCR-based method. DNA was extracted from representative colony of each isolate using the Kylt Purifier®, following manufacturer’s protocol. Typing was performed with a conventional in-house multiplex PCR. The resulting DNA fragments were separated and analysed using fully automated multicapillary electrophoresis system (QIAxcel system, Qiagen NV, Netherlands). DNA fragment sizes were determined relative to size standards and reference markers, and results were exported for electronic documentation. The final serotyping results presented are a consolidation of the two approaches. The PCR methods cannot distinguish serotype 1 from 14, and serotype 2 from 1/2 due to their highly similar *cps* loci, therefore, these are reported together.

### Antimicrobial susceptibility testing

The antimicrobial susceptibility of *S. suis* isolates was assessed using disc diffusion method as previously described [34]. Isolates were tested against a panel of antibiotics which were obtained from Oxoid™ (Thermo Scientific Sensititre, Waltham, MA, USA). The specific antimicrobials included in the testing panel were ampicillin (10 µg), amoxycillin + clavulanate (20/10 µg), penicillin (10 µg), ceftiofur (30 µg), cephalothin (30 µg), tetracycline (30 µg), sulfamethoxazole/trimethoprim (30 µg), and erythromycin (15 µg). Bacterial suspensions were prepared by selecting two to four colonies from freshly cultured, sheep blood agar plates. Colonies were resuspended in 5ml of demineralised water (Thermo Scientific Sensititre, Waltham, MA, USA), vortexed thoroughly and adjusted to turbidity equivalent to a 0.5 McFarland standard using densitometer (Serosep, Limerick, Ireland). The standardised inoculum was evenly spread onto Mueller-Hinton agar supplemented with 5% defibrinated sheep blood (Fannin L.I.P, Galway, Ireland) using a sterile swab. A maximum of four disks per 90 mm plate was applied to prevent zone overlap, following CLSI guidelines. Plates were incubated at 35°C in 5% CO_2_ for 20-24 hours. Zones of inhibition were measured and each testing batch included *Streptococcus pneumoniae* ATCC 49619 as a quality control strain. *S. suis* currently lacks species-specific clinical breakpoints in CLSI or EUCAST guidelines. Consequently, we used breakpoints (Table S2) established for other streptococcal species [35,36].

### Ethics statement

All diagnostic submissions were conducted as part of veterinary diagnostic services and farm health investigations; no live animals were experimentally infected for this study. Data use was in accordance with DAFM’s policy on research and diagnostic samples. All data were aggregated prior to reporting to safeguard farm-level confidentiality.

## Results

### Overview of submissions

Between 1st January 2010 and 31st December 2024, a total of 2,338 pig submissions comprising 4,921 pig carcasses were submitted to the VLS for post-mortem examination, as recorded in the LIMS database. Within this population, 150 submissions involving a total of 622 carcasses (12.6% of all carcasses) from 64 herds were examined. At least one *S. suis* isolate was archived from each of these submissions. In total, 170 *S. suis* isolates were archived from cases with a diagnosis of SSAD. Submissions originated from 21 counties across Ireland indicating a wide geographical distribution of SSAD. The number of carcasses per submission ranged from 1 to 10 (mean*=*4, median*=*3), indicating that practitioners often submitted multiple pigs when investigating outbreaks.

### Pathological findings in SSAD cases

Each case of SSAD was classified by the primary pathological diagnosis attributed to *S. suis* by the pathologist. In cases with multiple co-diagnoses, the most significant lesion was noted and used for classification. Consistent with *S. suis* associated disease, meningitis was the most common presentation, accounting for 41.8% (71 out of 170) of the submissions. A representative histological feature of meningitis is presented in Fig. 1, illustrating macrophage and neutrophilic infiltration within the meninges and adjacent the cerebral cortex consistent with acute suppurative inflammation.

**Fig. 1:**
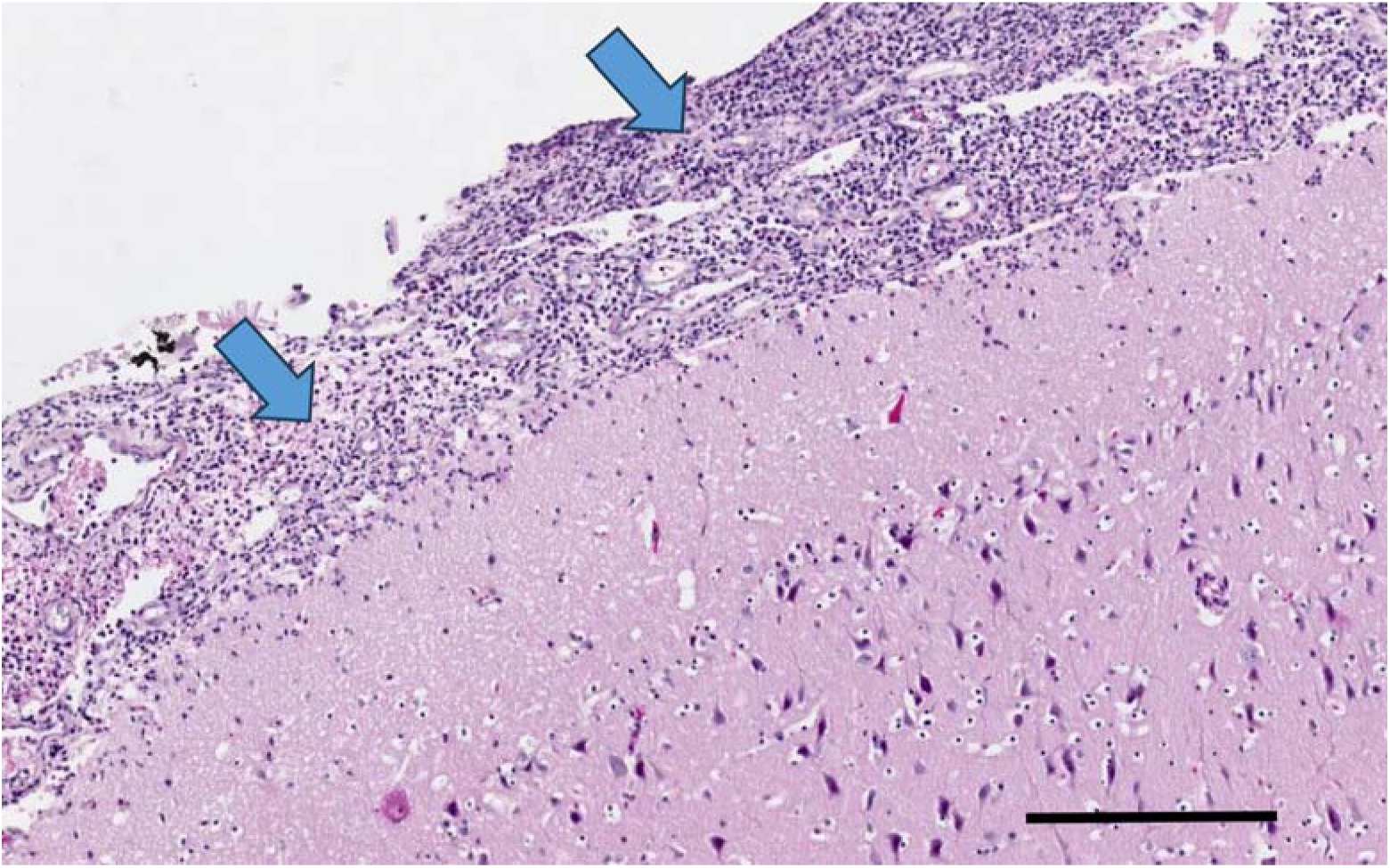
Representative histological section of a pig brain tissue showing acute meningitis. Haematoxylin and eosin-stained section shows dense infiltration of inflammatory cells (blue arrows), primarily neutrophils within the meninges, with inflammation extending into the adjacent cerebral cortex. Scale bar = 100µm

The next most common presentation was pneumonia (25.3%, *n*=43), followed by polyserositis (10%, *n*=17), arthritis (9.4%, *n*=16), bacteraemia/septicaemia (4.1%, *n*=7), endocarditis (2.4%, n=4), and pericarditis (2.4%, *n*=4) (Fig. 2). Additionally, *S. suis* was occasionally recovered from lesions which are not typically a hallmark of SSAD, including three cases of rhinitis, two cases of abortion, and a case of enteritis. Two submissions had no definitive lesion recorded despite yielding *S. suis* isolates.

**Fig. 2:**
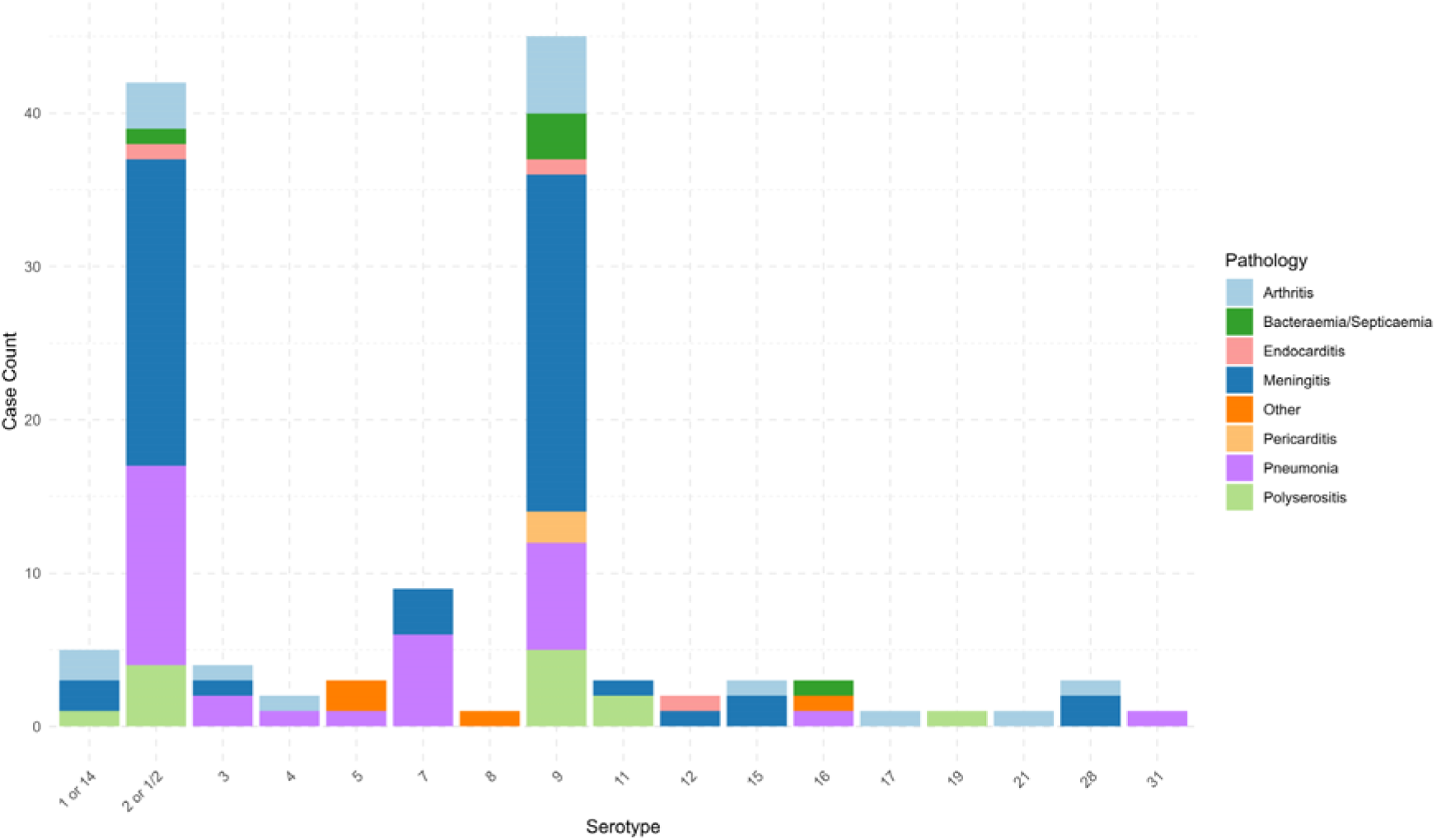
Distribution of SSAD pathological presentations by serotype. A stacked bar chart showing the number of cases (y -axis) of each serotype (x-axis) categorised by pathological diagnosis. Meningitis was the most common pathology recorded, followed by pneumonia, and polyserositis. Serotype 31 is included for completeness and is represented by a single isolate from a lung swab (non-carcass submission) in 2019.

### Serotype distribution of *S. suis* isolates

Of the 170 archived *S. suis* isolates from carcass submissions, 129 (75.9%) were successfully revived and serotyped. These originated from submissions representing 64 herds, each of which had at least one serotype identified. In total, 17 distinct *S. suis* serotypes, including serotype 31, were detected over the 14-year period (Fig. 3). Serotype 31 was represented by a single isolate obtained from a diagnostic lung swab (non-carcass submission) in 2019, increasing the total herd count to 65. The distribution was highly skewed toward a few dominant serotypes including serotype 9 and serotype 2 or 1/2, accounting for 35.7% (n=46) and 32.6% (n=42), respectively. The remaining one-third of isolates were distributed among 15 less common serotypes which individually accounted for less than 10% of the total, including serotypes 7, 1 or 14, 3, 4, 5, 8 11, 12, 15, 16, 17, 19, 21, 28 and 31. Specifically, only one isolate (<1%) was identified for serotypes 8, 17, 19, 21 and 31.

**Fig. 3:**
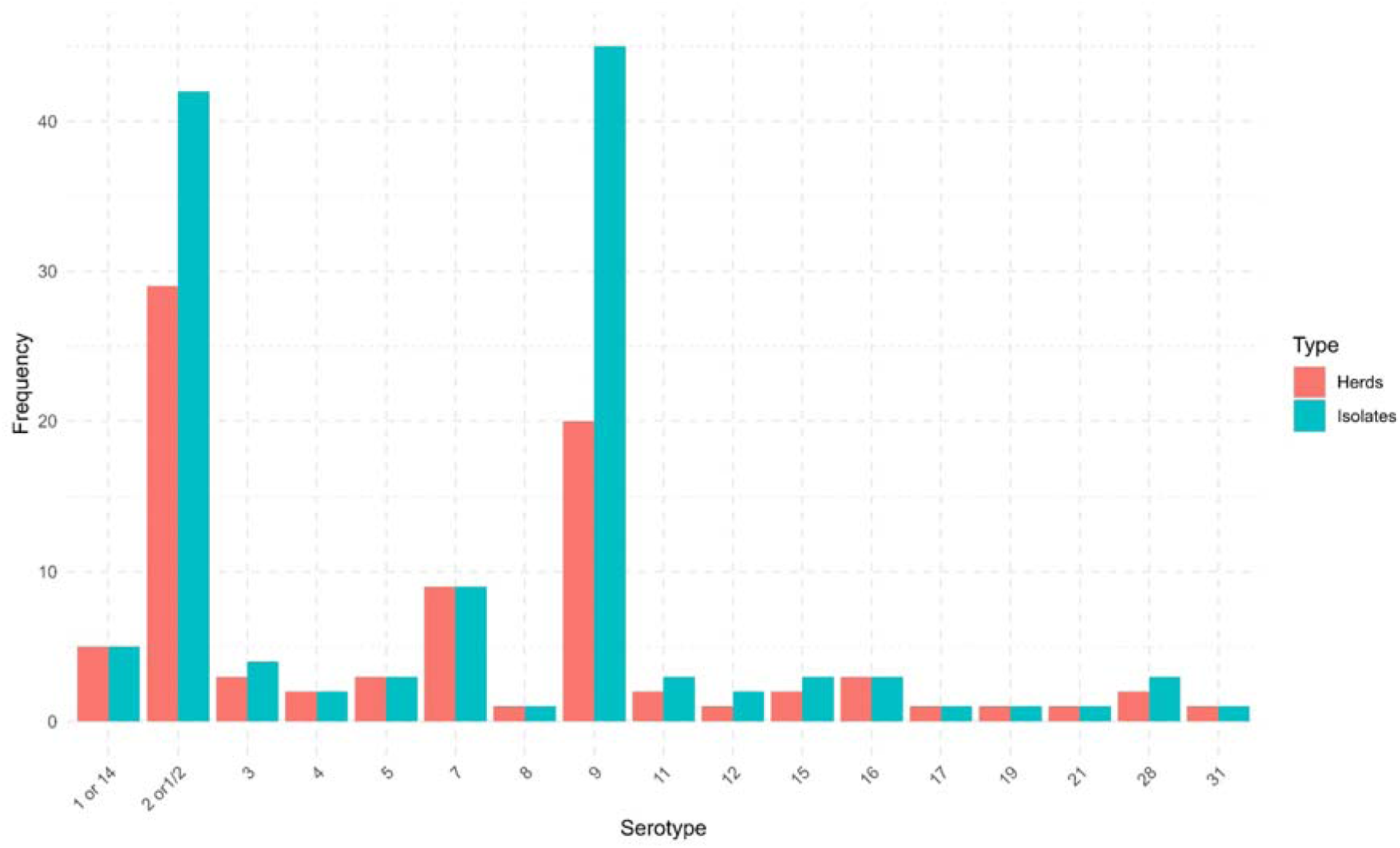
Distribution of serotypes among isolates and herds. A bar chart showing the frequency of 17 distinct S. suis serotypes among 129 archived isolates and 65 herds over a 14-year period. Serotype 2 or 1/2 and serotype 9 were the most frequently detected serotypes among all isolates. At the herd level, serotype 2 or 1/2 was detected in 44.6% of herds followed by serotype 9 (30.8%), and serotype 7 (13.8%). No other serotype was present in more than 10% of the affected herds.

At the herd level, serotype 2 or 1/2 was confirmed from carcasses in 29 of 65 herds (44.6%) that had detectable isolates, followed by serotype 9 (30.8%, *n=*20 herds) and serotype 7 (13.8%, *n=*9 herds). No other single serotype was present in more than 10% of the affected herds (Fig. 3). Most farms experienced a single dominant serotype causing outbreaks in that herd, although a few farms experienced multiple serotypes over time. It is noteworthy that the serotypes identified from SSAD carcasses encompass almost all those known to cause pig disease in Europe. We observed a marked increase in the proportion of serotype 9 isolates in later years from 2019 (particularly 2019–2024) (Fig. 4).

**Fig. 4:**
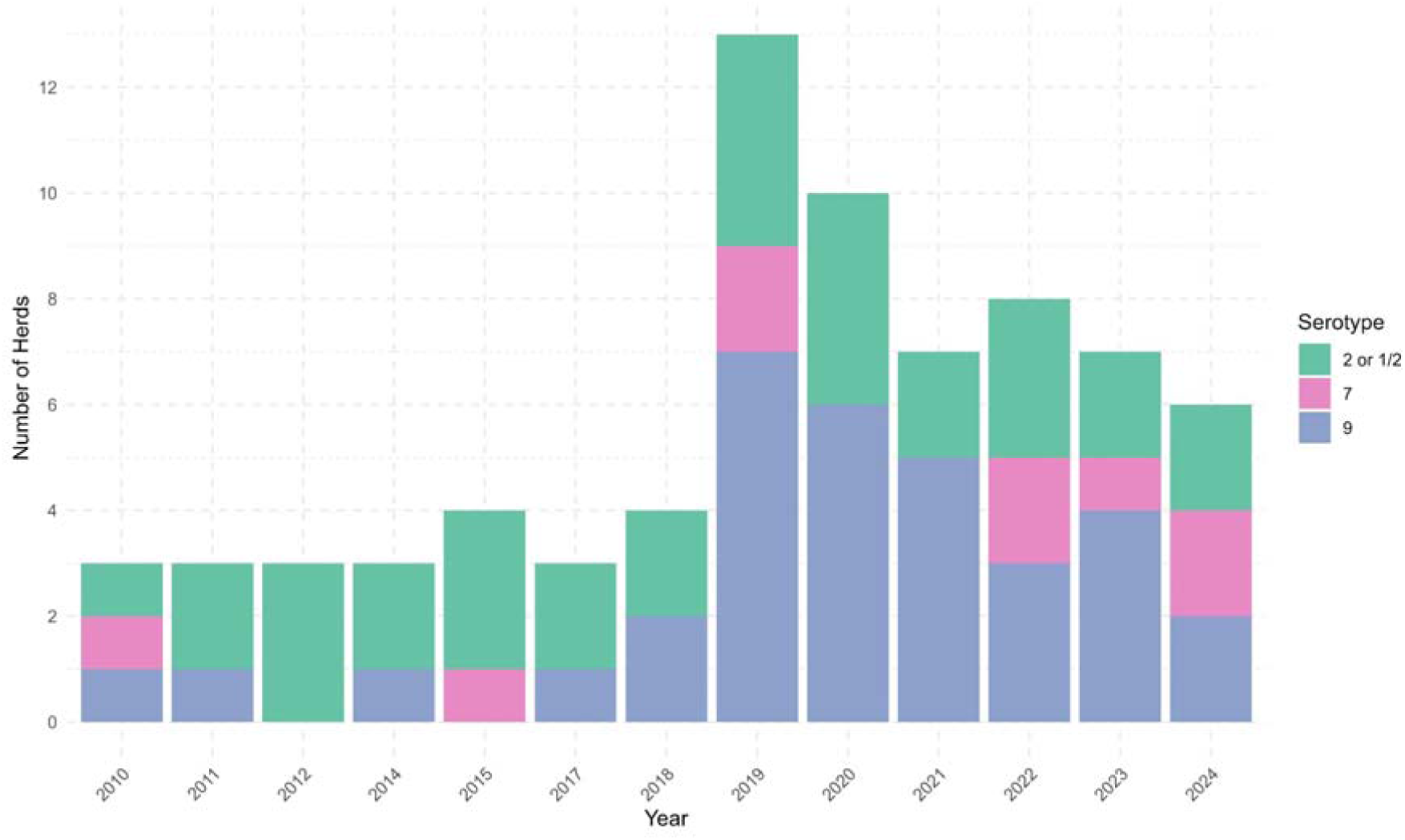
Temporal distribution of herds affected by serotypes 2 or 1/2, 7, and 9 between 2010 to 2024. An increase in the proportion of serotype 9 cases is evident in later years, particularly from 2019–2024. Conversely, serotype 2 or 1/2 remained relatively stable over time while serotype 7 appeared sporadically.

### Concurrent infections

Concurrent infections were frequently identified in pigs diagnosed with SSAD, highlighting the polymicrobial nature of many pig disease outbreaks, particularly those involving respiratory disease. In total, 48 animals with a confirmed diagnosis of SSAD had evidence of co-infection with at least one other significant viral or bacterial pathogen. For respiratory disease specifically, PCR testing for respiratory viral pathogens was performed on 33 of 43 pneumonia cases examined at necropsy. Of these, 22 (66.7%) tested positive for at least one viral pathogen. The most common viral co-infections in respiratory disease were: porcine reproductive and respiratory syndrome virus (PRRSV) in 9 of 28 animals tested (32.1%), swine influenza (SIV) in 6 of 27 (22.2%), and porcine circovirus type 2 (PCV 2) in 8 of 22 animals tested (36.6%). Across the entire dataset, PRRSV was detected in 18 (18%) out of 100 animals tested, SIV in 15 of 85 (17.6 %), and PCV2 in 18 out of 82 (21.9%).

In addition to *S. suis*, other bacterial pathogens were identified from lesions in some SSAD cases. *Actinobacillus pleuropneumonia* (APP), a primary cause of pleuropneumonia, was cultured from lung tissue in two cases of pneumonia and detected by PCR in cohort pigs of four other submissions. *Pasteurella multocida* was identified in one case, and co-existent *Pasteurella* pneumonia was identified in cohort pigs in another submission. *Glaesserella* (*Haemophilus) parasuis,* the agent of Glässer’s disease, was identified by PCR in three cases and in cohorts of three further submissions. *Mycoplasma hyopneumoniae* (enzootic pneumonia) was detected by PCR in four animals from each of four submissions and in cohorts from another submission. These are all pathogens that can co-exist with or predispose to *S. suis* infection, especially respiratory disease complex. In our dataset, fifteen herds experienced distinct outbreaks of polymicrobial pneumonia, with *S. suis* detected alongside one of the aforementioned bacteria.

### Antimicrobial susceptibility profiles of isolates

Between 2010 and 2024, *S. suis* isolates were tested for susceptibility to eight antimicrobials (Fig. 5). The number of isolates tested for each antimicrobial varied due to differences in panel composition and reagent availability over the years, with testing becoming more standardised and comprehensive over time. Resistance patterns differed considerably across antimicrobials. Tetracycline resistance was widespread, observed in 66% (98/149) of isolates. Resistance to tetracycline increased gradually over time, rising from 66.7% in 2010 and 2017 to over 70% from 2019 onwards, and peaking at 81% in 2022 before dropping to 69.6% in 2023. Erythromycin resistance was detected in 44.4% (64/144) of isolates and remained relatively stable, with slight fluctuations between 35% and 54% from 2018 onwards. Sulfamethoxazole/trimethoprim resistance was present in 23.5% (12/51) of isolates. Conversely, resistance to penicillin (7.89%, 12/152) and ampicillin (5.5%, 9/164) remained low throughout this study. Throughout the study period, the isolates consistently maintained low resistance levels to these beta-lactam antibiotics. Cephalothin resistance was equally low (3.3%, 5/151), while ceftiofur resistance was rare (1.4%, 2/145). Interestingly, no isolate (0/157) demonstrated amoxycillin/clavulanate resistance throughout the period.

**Fig. 5:**
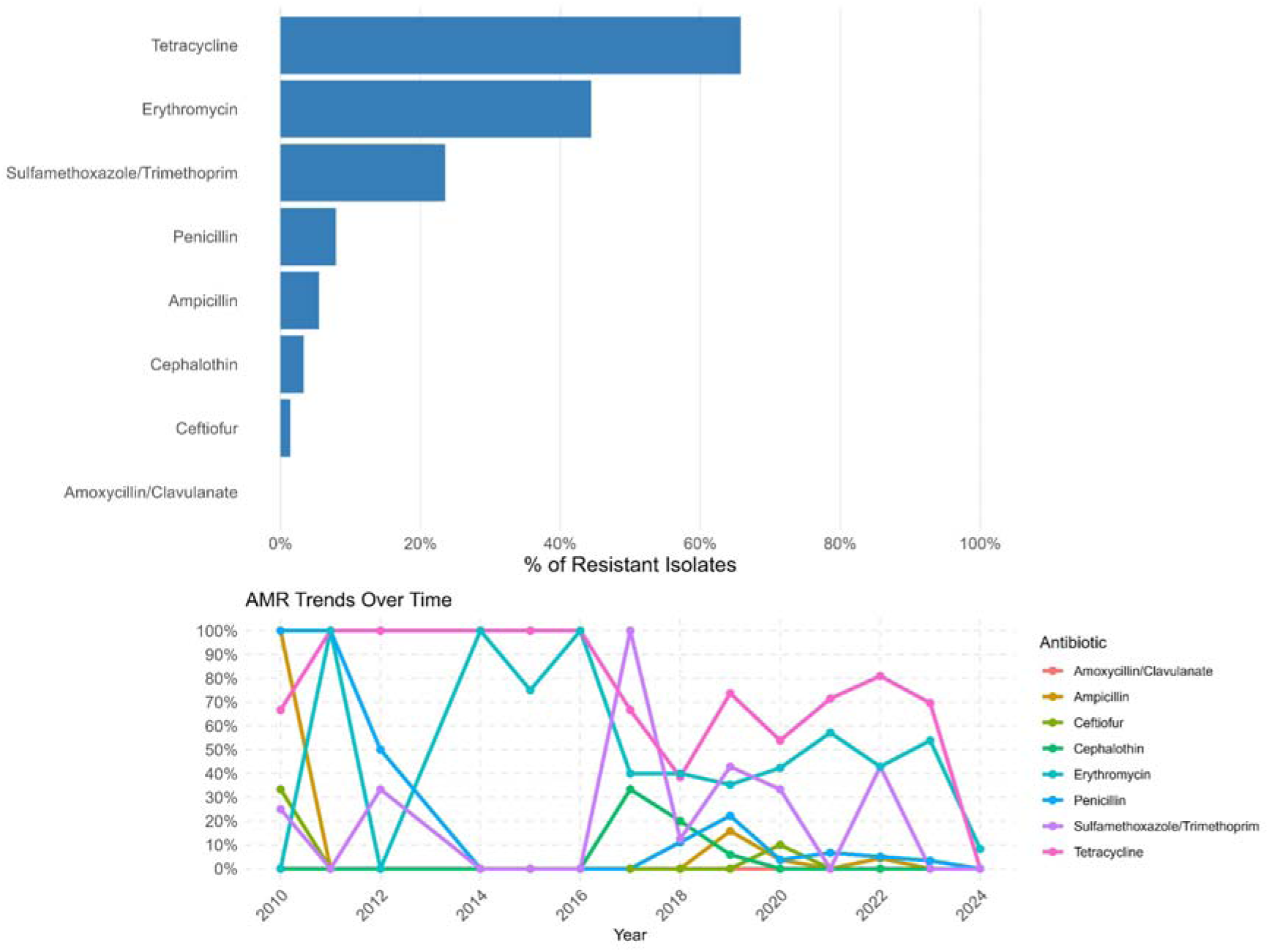
Antimicrobial resistance profiles of S. suis isolates over time. Bar chart (top) shows the overall percentage of resistant S. suis isolates for each of the 8 antibiotics tested. Line plot (bottom) illustrates yearly resistance trends over time. Resistance was highest for tetracycline. Resistance to beta-lactams (penicillin, amoxycillin/clavulanate, and ampicillin) remained low or absent throughout the study period.

## Discussion

This study provides the first comprehensive overview of SSAD in the Irish pig population, involving 14 years of diagnostic cases. Approximately 7.6% of all pig diagnostic submissions to the state laboratory service were attributed to SSAD, indicating a significant baseline level of *S. suis* impact on the national herd. This figure also demonstrates that *S. suis* remains a major cause of disease and losses in Irish pig farms, alongside other endemic diseases. Clinically and pathologically, the disease manifestations are consistent with descriptions of SSAD in the literature. Meningitis was the leading presentation in our dataset (Fig. 2), which is consistent with reports that serotype 2 classically causes meningitis and polyarthritis in weaned pigs. Several cases were associated with pneumonia or polyserositis, conditions often associated with *S. suis* either as a primary agent or in conjunction with other pathogens. Moreover, we recorded a few atypical presentations, such as *S. suis* isolated from abortion and enteritis cases. These could represent an instance of *S. suis* acting as an opportunist or secondary invader; however, their rarity suggests that *S. suis* should mainly be considered a pathogen of the nervous system, serosal cavities, and joints of pigs, and in lung, often as part of the porcine respiratory disease complex (PRDC).

We identified 17 distinct serotypes among 129 isolates, including a single serotype 31 isolate obtained from a lung swab. Whole-genome sequencing confirmed the isolate as virulent. Although serotype 31 is rarely reported, its detection expands the range of pathogenic serotypes observed in Ireland. This substantial serotype diversity is comparable to reports of other European countries with molecular typing programmes including Switzerland [37,38]. Segura *et al.* noted that while numerous serotypes are implicated in disease globally, only a few tend to dominate [17]. In our data, serotype 9 and serotype 2 or 1/2 dominated, accounting for over two-thirds of all cases. This observation is consistent with recent trends in Europe over the past two decades in which serotype 2 or 1/2 was historically the most virulent and prevalent. However, in Europe, serotype 9 has emerged as a leading cause of pig disease since 2010s [17]. Our findings reveal a similar shift in Ireland. Since 2019, serotype 9 has become increasingly prominent in Irish SSAD cases, surpassing serotype 2 or 1/2, which is apparent in both carcass submissions and at herd level between 2019 and 2024 (Fig. 4). While the reason behind this shift is not fully understood, several hypotheses have been proposed. One possibility for the increased occurrences of serotype 9 is the rapid expansion and globalisation of the pig industry, particularly via the international movement of seed stock, which may have facilitated the dissemination of successful serotype 9 clones [39,40]. Another hypothesis explores vaccine-driven serotype replacement. In regions where autogenous or commercial vaccines targeting serotype 2 have been administered extensively, they may suppress the circulation of serotype 2 and create an ecological space for other serotypes, such as serotype 9 to become more prevalent [23]. Additionally, other contributory factors may include natural evolutionary advantages or unknown virulence-associated factors that favour the success of serotype 9.

Vaccination is an essential tool in *S. suis* management on Irish pig farms. Where commercial vaccines are not available for a given serotype, autogenous vaccines are often used [16]. The autogenous vaccines are produced from *S. suis* strains isolated from clinically affected animals and are prepared by specialist laboratories for use exclusively on the originating farm. Such vaccines can be highly effective in herds where a single serotype remains dominant over time and the virus status within the herd is stable. However, the lack of cross-protection between serotypes presents a major challenge in vaccine implementation [41]. Our findings, along with other studies, show that the dominant serotype in herds can change, or multiple pathogenic serotypes can circulate and cause disease at different times. In such cases, an autogenous vaccine formulated from previous seasons may provide limited protection if new serotypes emerge. These dynamics necessitate continuous on-farm surveillance, even where vaccination protocols are established. From a diagnostic perspective, relying on a single isolate from a necropsied pig may not adequately represent an entire outbreak, particularly in herds with documented serotype variability. Higgins and Gottschalk [42] recommended collecting two or more alpha-haemolytic colonies from different tissues of the same animal or different animals within the same cohort. Our data supports this approach. Farms with varying SSAD lesions across animals or recurring outbreaks despite vaccination would merit broader sampling for isolates to identify emerging or co-circulating pathogenic serotypes.

The frequent detection of concurrent infections in SSAD respiratory cases in this study aligns with the growing consensus that *S. suis* is often part of a disease complex rather than a standalone pathogen. Respiratory viruses are known to compromise lung defences and modulate the immune system, creating an opportunity for *S. suis* to invade the bloodstream and tissues. For instance, a recent Spanish case-controlled study found that PRRSV infection at weaning increased the odds of *S. suis* disease by a factor of 6.7 [6]. Beyond viruses, we detected concurrent bacterial infection in several cases, especially APP and *G. parasuis*. These mixed infections can complicate both diagnosis and treatment. The presence of concurrent infections means veterinarians must consider integrated, broad-spectrum strategies and address underlying management issues such as ventilation, stocking density, or herd immunity gaps to break the cycle of disease [21,43]. Maintaining effective vaccination programmes for concurrent infections on farm and robust biosecurity measures may indirectly help control *S. suis* losses.

We observed very low resistance levels to beta-lactam antibiotics including penicillin, ampicillin, amoxycillin/clavulanate, and ceftiofur (Fig. 5). Beta-lactams, especially penicillin or amoxycillin, are first-line treatment for *S. suis* in pigs, and their continued effectiveness is important to both production costs and One Health. Our results indicate that, as of 2024, Irish isolates remain broadly susceptible to these drugs, which is consistent with surveys from other European countries. Dechêne-Tempier *et al.* [44] reported that in France, only 5% of strains showed resistance to beta-lactams. Similar levels of susceptibility to beta-lactams have been reported in Switzerland, the Netherlands, and Czechia [45–47]. In contrast, high resistance of *S. suis* to tetracyclines and macrolides is well-founded in global studies [25]. Resistance to tetracycline was observed in two-thirds of the Irish isolates. While significant, this rate is lower than the proportion reported in Denmark (75%), France (81%), England (91%) and North America (95.5%) and China (97.8%) [44,48–50]. Macrolide resistance among the Irish isolates was lower than has been reported in Denmark (61%) and France (54–60%) [44,51]. Overall, the high susceptibility to beta-lactams, despite their use in pigs, is a positive finding, and these antibiotics remain the recommended first-line therapy.

The Irish pig industry is a major contributor to national agricultural output (behind beef and dairy). *S. suis* outbreaks carry significant economic overheads, including piglet mortality, increased treatment costs, labour demands, and potential welfare implications. Furthermore, effective control also supports national efforts to reduce antibiotic use by preventing disease occurrence through vaccination and appropriate disease management practices, thereby reducing reliance on antimicrobials. This baseline data is therefore essential for informing targeted, evidence-based control strategies. It is important to note that these findings are based on ad hoc diagnostic submissions, as such, the patterns described here reflect the characteristics of diagnosed cases rather than the overall prevalence of SSAD in the national herd. Nonetheless, these data are valuable for understanding the relative importance of these serotypes in disease occurrence and for guiding control efforts.

Future studies should investigate the molecular epidemiology of the Irish isolates to assess genetic relationships with global lineages. In particular, it would be useful to assess whether the Irish serotype 9 isolates are related to expanding lineages including ST1 and ST16, which have been associated with this serotype in other European countries [38]. Whole-genome sequencing of archived isolates may reveal novel resistance or virulence-associated factors. Moreover, development of *S. suis*-specific breakpoints for antimicrobial susceptibility testing across a broader range of antibiotics is needed, particularly in settings without routine access to molecular typing and whole-genome sequencing. Additionally, data on efficacy of autogenous vaccines in the field and more formally through trials are also needed, as current information is limited.

### Conclusion

This study has generated baseline data on *Streptococcus suis*-associated disease in Ireland. Serotype 9 was the most frequently identified overall, driven by a notable increase in its occurrence since 2019, surpassing serotype 2 or 1/2 in SSAD in Ireland. Although control of *S. suis* is complicated by co-infections and resistance to some antimicrobials, susceptibility to penicillin and amoxycillin/clavulanate, the first-line therapy, remains high. Going forward, evidence-based integrated approaches including improved on-farm biosecurity to reduce stress and co-infections, implementing serotype-specific vaccination where appropriate, and maintaining prudent antibiotic use, are essential. Such measures, supported by ongoing surveillance, will help mitigate the impact of SSAD and improve the health and productivity of the Irish national pig herd.

## Availability of data and materials

Datasets used or analysed during the current study are available from corresponding author on reasonable request. Primers information and antibiotic breakpoints are provided in supplementary tables (Table S1 and Table S2).

## Acknowledgements

Authors would like to thank all the pig farmers and their specialist pig veterinary practitioners for providing submissions for diagnosis, information and assistance. We are also grateful to the DAFM staff in the Central and Regional Veterinary Laboratories for collecting data, performing post-mortem examinations, and conducting sample testing. Particular thanks to Kevin Kenny for suggestions on the manuscript.

## Author contributions

**John Moriarty:** Writing – original draft, review & editing, supervision, methodology, investigation, funding acquisition, formal analysis, data curation, conceptualisation. **Emmanuel Kuffour Osei:** Writing – review & editing, software, methodology, formal analysis, data curation, conceptualisation. **Colm Brady:** Writing – review & editing, methodology, investigation, formal analysis, data curation. **Sara Salgado:** Writing – review & editing, methodology, investigation, formal analysis, data curation. **Sebastian Alessandro Mignacca:** Writing – review & editing, investigation, formal analysis, data curation. **Fiona Kane:** Writing – review & editing, methodology, investigation, formal analysis, data curation. **Dayle Johnston:** Writing – review & editing, methodology, investigation, formal analysis, data curation. **Laura Scanlon:** Writing – review & editing, methodology, investigation, formal analysis, data curation. **Hannah Flynn:** Writing – review & editing, methodology, investigation, formal analysis, data curation. **Marius Beumer:** Writing – review & editing, methodology, investigation, data curation. **Rachel Reid:** Writing – review & editing, methodology, investigation, formal analysis, data curation. **Philip Curran:** Writing – review & editing, methodology, investigation, formal analysis, data curation. **Jennifer Mahony:** Writing – review & editing, methodology, investigation, formal analysis, data curation. **Aine O’Doherty:** Writing – review & editing, methodology, investigation, formal analysis, data curation. **Shane McGettrick:** Writing – review & editing, methodology, investigation, formal analysis, data curation. **Cosme Sánchez-Miguel:** Writing – review & editing, software, methodology, investigation, data curation. **Margaret Wilson:** Writing – review & editing, methodology, investigation, formal analysis, data curation. **John G. Kenny:** Writing – review & editing, methodology, investigation, funding acquisition, formal analysis, data curation.

## Consent for publication

Each author contributed to this work and gave final approval of the version for publication.

